# Contextual Cues and Transition Statistics Drive Expression of Competing Motor Memories

**DOI:** 10.1101/2025.09.08.674705

**Authors:** Adarsh Kumar, S Adith Deva Kumar, Sumit Sannamath, Neeraj Kumar

**Author notes:** These authors contributed equally to this work (Adarsh Kumar, S Adith Deva Kumar). DATA AVAILABILITY: The datasets used and/or analyzed during the current study are available from the corresponding author upon reasonable request.

## Abstract

Learning multiple motor skills without interference and expressing the correct one in a changing environment is a fundamental challenge. Contextual cues are known to help separate these memories, but how they interact during retrieval is not well understood. We investigated how the stability, recency, and transitional statistics of learning environments influence this process. Across six visuomotor adaptation experiments, participants learned opposing rotations (Tasks A and B) tagged with distinct contextual cues under different schedules (blocked or interleaved) and were tested in stable or dynamic environments. We found that while contextual cues can successfully separate memories, expression is systematically biased by learned transition statistics: towards more stable memories after imbalanced training, and towards more recent memories when stabilities are matched. Critically, when the stable statistics of training mismatched the volatile statistics of testing, cue-based retrieval collapsed, and behavior was dominated by these stability or recency biases. Conversely, learning in a high-entropy, interleaved environment enabled precise, cue-appropriate expression regardless of the testing schedule. These results demonstrate that memory retrieval is not cue-driven but arises from an arbitration process between cues and transition priors. Our findings reveal that memory retrieval involves weighting sensory information against latent priors derived from the history of context transitions. This work provides a unifying theoretical framework for understanding adaptive memory expression, positing that the brain leverages the learned statistical structure of the environment to infer which memory to recall, thereby balancing cue-driven selection with the stability and predictability of past experience. This principle offers a unifying explanation for interference, spontaneous recovery, and the benefits of variable practice, providing a more holistic model of adaptive motor behavior.

**Statement of Significance:** How does a tennis player instantly switch between a forehand and a backhand? Our work reveals a fundamental principle of how the brain organizes and retrieves memories. We demonstrate that recalling a skill is not just about recognizing a contextual cue, but about an internal process of integrating that cue with the learned statistics of the environment, such as the stability and recency of past experiences. This finding provides a unifying framework for phenomena like interference and spontaneous recovery. It has significant implications for designing more effective training in sports and rehabilitation, where structuring practice around environmental statistics can optimize learning and promote flexible skill application.

## Introduction

The brain’s ability to select and express the right action from a vast repertoire of learned behaviours is a cornerstone of adaptive behaviour. Consider a tennis player preparing to return a serve, a chef switching between knives, or a musician adapting to different instruments. Years of practice hone distinct motor plans for each context, yet in the fleeting moment of action, the brain must swiftly suppress an anticipated response if the context shifts. This challenge of retrieving the correct memory amidst uncertainty extends beyond motor tasks to episodic, working, and language memory, highlighting a fundamental question in memory research: how does the brain retrieve the correct memory from a library of learned representations, particularly when contexts are ambiguous or rapidly changing?

This retrieval problem is pronounced when multiple memories compete for expression. Performance errors often stem not from forgotten skills but from expressing the wrong memory. In motor learning, studies of visuomotor adaptation reveal this distinction: when participants learn opposing perturbations (e.g., clockwise and counterclockwise rotations) in sequence, they exhibit anterograde interference, struggling to re-express the first skill after learning the second, despite intact initial learning^1,2^. These errors reflect retrieval failures, not acquisition deficits, underscoring the need for mechanisms that guide memory selection.

Traditional theories propose that contextual cues, such as specific postures, workspaces, or visual stimuli, solve this problem by tagging memories for later retrieval^3^. For example, associating a motor skill with a unique cue enables rapid recall, as seen in savings, where re-encountering a perturbation elicits faster relearning^4^. However, cues have limitations. Many are weak, ambiguous, or overlapping in real-world settings, and their effects are graded rather than absolute. For instance, varying premovement cues along a continuum results in a weighted blend of associated memories, consistent with Gaussian generalization^5^. Moreover, retrieval can fail when cue-context relationships are disrupted, such as when blocked training with visual cues is followed by interleaved testing, leading to increased interference and impaired memory separation across domains^6,7^. These findings suggest that cues alone are insufficient for robust memory retrieval in dynamic environments.

We propose that the brain relies on transition statistics - learned expectations about how contexts evolve over time; as a fundamental mechanism for guiding memory retrieval across domains. Transition statistics encompass volatility (how frequently contexts change) and sequential regularities (which contexts follow others). For example, if a learner expects context A to typically precede context B, being in A primes anticipation of B, facilitating retrieval of the associated memory. In stable environments with rare switches, the brain persists with the current memory; in volatile settings with frequent changes, it remains flexible, ready to switch. This framework unifies memory organization across motor control, episodic, working, and language domains, offering a novel principle for adaptive behavior.

Evidence for transition statistics spans multiple domains. In motor adaptation, learning rates adjust to environmental volatility: stable perturbations accelerate adaptation, while unpredictable switches slow it, consistent with Bayesian learners downweighing errors in high-volatility settings^8^. Similarly, cues signalling context repetition enhance single-trial adaptation, while cues indicating change dampen it^9–11^. In skill acquisition and procedural memory, implicit sequence learning tasks, such as the serial and alternating serial response time tasks, shows sensitivity to high-probability motor transitions even without conscious awareness^12,13^. In episodic memory, abrupt context shifts (event boundaries) segment experiences, reducing interference, while stable contexts promote integration^14,15^. In spatial memory, the hippocampus encodes probabilistic transition graphs between locations, enabling flexible route planning and also supports cognitive map formation^16^. Infants leverage transitional probabilities to segment speech into words, detecting structure without explicit cues^17,18^. Bilingual individuals who frequently switch languages exhibit smaller switch costs, reflecting flexible retrieval shaped by transition experience^19^. In the attention domain, probabilistic cueing and temporal expectation paradigms demonstrate that allocation of attention is guided by the learned transition statistics of where and when events are likely to occur^20,21^. In perceptual decision-making, humans adapt cautiously in volatile environments, guided by inferred transition structures^22^. In working memory, temporal clustering of items influences recall order, driven by learned transition probabilities^23^. These parallels suggest that transition statistics are a domain-general mechanism for organizing memory retrieval. Brain learns not just memory-cue associations but also the temporal structure of contexts, enabling a meta-learning process that optimizes retrieval across motor, episodic, and other memory systems.

Mechanistically, this process is formalized through Bayesian context-inference models, such as the Contextual Inference (COIN) model^9–11^. Context is a latent variable, and the brain maintains a probabilistic belief over possible contexts, updated via sensory cues, recent errors, and transition priors. Retrieval reflects a posterior distribution combining these factors, not just cue-driven selection. High volatility shifts expression toward recent memories, while stability favours more experienced ones. This process aligns with neural mechanisms where motor cortex populations simultaneously encode multiple action plans, weighted by context likelihood^24^. The principle of transition statistics extends to neural population codes, where dynamic factor combinations encode context-dependent memories^25^.

To test this theory, we conducted six visuomotor adaptation experiments, manipulating cue presence, training order, stability, and transition entropy. Participants learned opposing motor mappings with distinct cues under blocked or interleaved schedules, then expressed them under varied cue and transition conditions. Our findings reveal how transition statistics shape memory expression, offering behavioural evidence for a domain-general framework with implications for neural mechanisms, skill learning, and rehabilitation.

## Methods

### Participants

A total of 144 healthy right-handed participants (81 men, 63 women, age [mean ± SD] = 22.24 ± 2.6 years, 12 individuals per experimental group) with normal or corrected-to-normal vision participated in the study. The participants reported no neurological disorders, cognitive impairments, or orthopedic injuries. The Edinburgh Handedness Inventory was used to evaluate the handedness of the participants^26^. All participants gave written informed consent and were naive to the purpose of the study. On successful completion of the task, participants were monetarily compensated for their time. The Institute Ethics Committee approved the study.

### Experimental setup

The participants sat in a dark and sound-attenuated room on a height-adjustable chair in front of a virtual reality frame setup on which a digitizing tablet was placed (GTCO CalComp). Participants made planar reaching movements on the tablet using a stylus. A high-definition display screen (1920 x 1080 Pixels, 120 Hz) was mounted on top and projected downwards onto a semi-transparent mirror. The mirror was placed between the screen and tablet, which occluded participants’ vision of their arms, and they had to rely on the cursor projected onto the mirror from the screen to get indirect visual feedback of their hand (Figure 1a). The hand position was recorded at a sampling rate of 120 Hz by the digitizing tablet.

**Figure 1:**
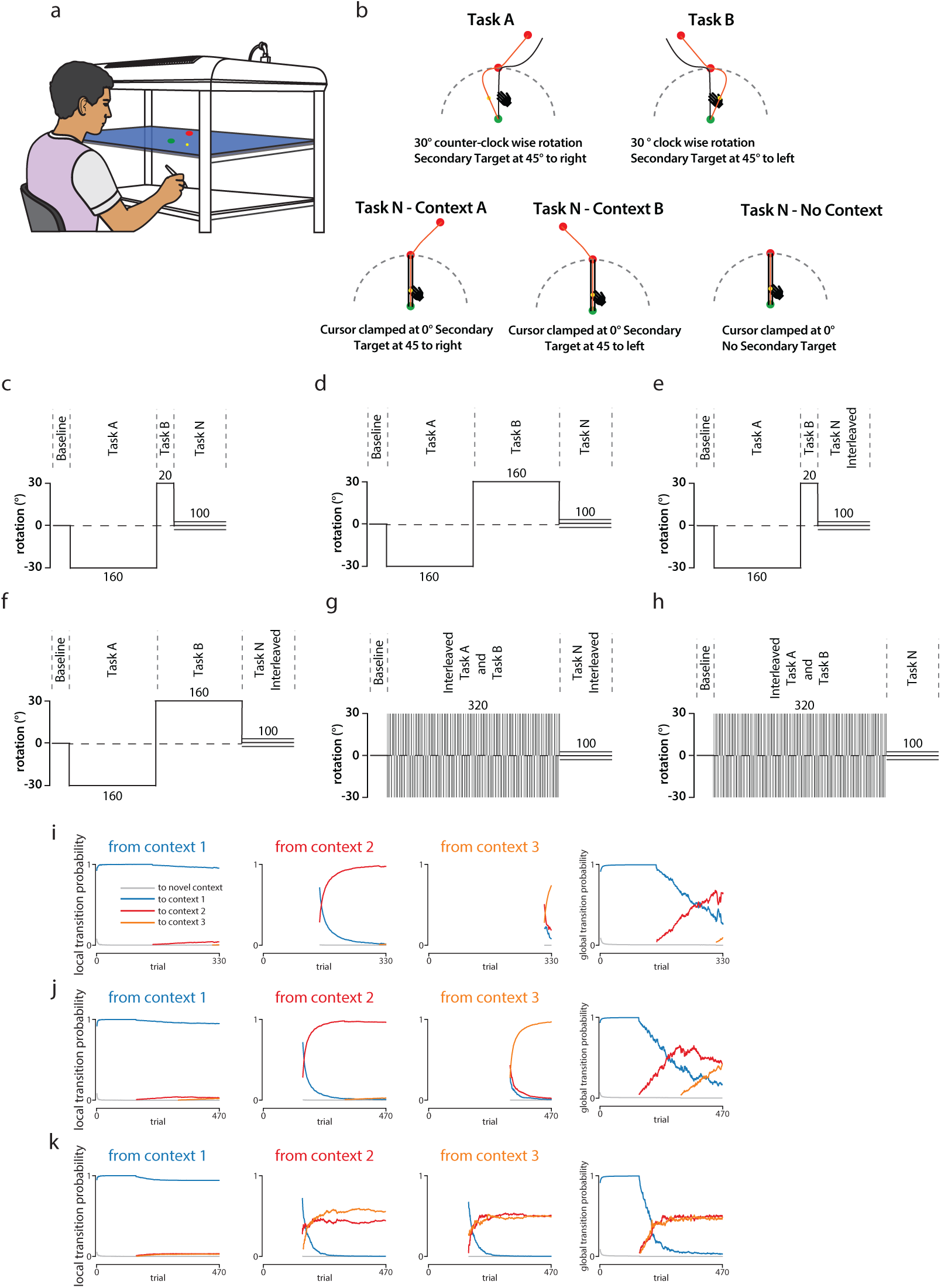
Experimental Procedure and learning schedules. (a) Sketch of experimental setup used to present start and targets. (b) Motor learning tasks. During memory acquisition, Task A (clockwise visuomotor rotation) and Task B (counter-clockwise visuomotor rotation) had opposite perturbations. The secondary target was a cue attached to the visuomotor rotation applied during learning. Memory expression had task N (zero-degree error clamped visuomotor rotation) and had one of the cues of Task A and Task B, or no cue. All the perturbations were only applied for the primary reach. The secondary reach didn’t have any imposed perturbation. The red line indicates the cursor trajectory, and the black line indicates the hand path. The hand symbol depicts instantaneous hand position, and the yellow dot represents the rotated cursor position on the screen. In Experiment 1, three independent groups of participants were exposed to either cue A, cue B, or no cue during Task N to test interference effects between memories formed in Task A and Task B. Experiment 1a (panel c) assessed stability bias (longer Task A than Task B). In contrast, Experiment 1b (panel d) examined recency bias (equal Task A and Task B). Experiment 2 introduced dynamic presentation of all three cues during Task N, with Experiment 2a (panel e) testing performance after a short Task B and Experiment 2b (panel f) after a long Task B. In Experiment 3, perturbations from Tasks A and B were interleaved during learning, with expression tested during Task N under two conditions: Experiment 3a (panel g) dynamically presented all three cues, whereas Experiment 3b (panel h) tested each cue in separate participant groups.(i, j and k) local and global transitional probability for each learning schedule.

The hand-controlled yellow cursor (0.3 cm diameter) indicated the stylus position. Each trial began when the cursor entered a blue start circle (0.8 cm diameter). After 550 milliseconds, the start target turned green, marking it as a go signal for the participants to initiate the reach to the primary and secondary targets (red colour, diameter = 0.8 cm) in sequence. Participants were required first to reach the primary target (straight in front of the participant, distance -12 cm) and then to the secondary target located 30° to the right (contextual cue A) or to the left (contextual cue B) at the same radial distance from the primary target. The primary reach was treated as the main task, while the secondary reach served as a contextual cue (Figure 1b).

After familiarization with the setup, instructions for the task, and a few practice trials, participants were asked to perform three consistent blocks: a baseline block (150 trials), a memory acquisition block (variable number of trials depending on the experiment), and a memory expression block (100 trials). In the baseline block, the cursor feedback was veridical to the hand movement, and it established unperturbed motor behavior and familiarized participants with the trial structure. For the first 50 trials, the secondary target was not presented, and participants had only to make the primary reach. For the subsequent 100 trials, one of the two secondary targets was also presented randomly.

Following the baseline session, the memory acquisition block was introduced, where the cursor movement was rotated relative to the hand movement during the primary reach. After the cursor reached the primary target, the rotation was removed, and the cursor moved in the same direction as the hand for the secondary reach. The location of the secondary target was linked with the direction of the perturbation imposed during primary reach. If the second target was presented on a clockwise/right side (cue A), then the perturbation for primary reach was a 30° counterclockwise cursor rotation (Task A). If it was presented on a counterclockwise/left side (cue B), then the cursor rotation for primary reach was 30° clockwise (Task B). This way, the secondary target acted as an explicit contextual cue about the upcoming perturbation in each trial. The number and order of these perturbation trials during the acquisition block varied across groups. The acquisition block was followed by a memory expression block where the cursor movement from the primary reach was clamped at 0° to the target, and the movement feedback to the secondary target from the first was veridical. The secondary target was presented either on the right for context A, on the left for context B, the same as the memory acquisition block, or was absent for no context trials. Participants were not notified about the change in the block (Figure 1b).

At the end of each trial, the participant was provided feedback about performance and speed. A numerical score represented the performance/reward about the reach where 10 points were given if the cursor ended within the target circle at the end of the movement, 5 points if the cursor was within 0.25 cm from the target edge, 1 point if within 0.4 cm and 0 points was provided if the endpoint of the cursor was beyond this distance. The overall score was also presented with the cumulative sum of the points obtained. Participants were asked to maximize the score. Participants also received text on screen as “Too slow” in red if their peak velocity was less than 32 cm/s, “Good” in green if it was between 32-50 cm/s, and “Too Fast” in red if the peak velocity was above 50 cm/s. The points obtained did not impact the compensation provided to the participants at the end of the experiment, and the points were also not analyzed. The performance feedback was not provided during the memory expression block.

### Experiment 1

The first experiment was designed to investigate whether the stability of acquired memory would affect how the memories are expressed. We hypothesized that a more stable context with low transitional probability would have a higher chance of being expressed. Furthermore, when multiple memories are equally stable, we expect the most recent context to influence expression more.

#### Experiment 1a

In our first experiment (1a, N=36), we first aimed to understand whether contextual cues can prevent interference even if one of the memories is acquired for a very short time. If yes, would the expression of memories be biased by the stability of contextual memories? Participants were randomly assigned to three groups that differed in the contextual cue presented during the memory expression block. After the baseline block, participants learned Task A (with cue A) first for 160 trials, followed by Task B (with cue B) for only 20 trials (Figure 1c). In three separate groups (N=12 per group), participants either received cue A, B, or no secondary target during the memory expression block.

#### Experiment 1b

Experiment 1b (N=36) aimed to determine whether the most recent memory would be expressed if multiple memories were equally stable. The experimental groups were similar to the first experiment, except that Task B was learned for the same number of trials as Task A (160 trials) during the acquisition block (Figure 1d).

### Experiment 2

We next investigated whether attaching distinct contextual cues to memories in separate sessions would be sufficient for distinct contextual expression in a dynamic, highly volatile environment and whether the stability of contextual memories would help prevent interference.

#### Experiment 2a

In Experiment 2a (N=12), we investigated whether attaching contextual cues to competing memories where one memory is only learned for a very short period, like Experiment 1a, would prevent interference if expressed in a randomly changing environment. To test this, a group of participants learned Task A and B, similar to experiment 1, but were presented with cue A, cue B, or no cue trials in a randomly interleaved manner during the expression block (Figure 1e). Thus, while they learned the tasks in blocked stable sessions, the expression was in a dynamically changing environment.

#### Experiment 2b

In this experiment (N=12), we asked if the interference observed in experiment 2a was because Task B was only learned for a limited time compared to Task A. To test this, a group of participants learned task A and task B, like experiment 1b. However, they were presented with cue A, cue B, and no cue trials randomly interleaved, as in experiment 2a (Figure 1f).

### Experiment 3

The third experiment was designed to investigate whether acquiring multiple contextual memories in a high transitional probability environment would be necessary to tag memories so they can be expressed distinctly in dynamic and stable environments.

#### Experiment 3a

In Experiment 3a (N=12), we aimed to investigate whether contextual cues could prevent interference if multiple memories are learned in a dynamically changing environment and expressed in a similar environment. To test this, a group of participants learned both Task A and Task B interleaved during the memory acquisition block, and the memory expression block included all three contextual conditions presented randomly in Task N. The Experiment consisted of 150 baseline trials, 320 interleaved memory acquisition trials, and 100 interleaved memory expression trials with cue A, B, or no cue (Figure 1g).

#### Experiment 3b

In Experiment 3b (N=36), we aimed to unravel whether multiple memories can be expressed in a constant environment if learned in a dynamically changing environment. In this experiment, participants were randomly assigned to 3 groups that varied with the contextual condition presented during Task N in the memory expression block. After 150 baseline trials, participants were presented with 320 randomly interleaved Task A and Task B trials. Participants were expected to simultaneously learn both visuomotor rotations in this block with their respective contextual cues. The memory expression block consisted of 100 clamped trials where one of the contextual conditions was presented in Task N according to their assigned group (Figure 1h).

### Data analysis

The data obtained were analyzed using custom MATLAB scripts. The hand position data were filtered using a low-pass Butterworth filter with a 10 Hz cut-off frequency. The speed of movement was calculated by differentiating the position data. The moment at which hand speed initially exceeded 5% of maximum movement speed was identified as the movement onset. The direction error was calculated as the angle between the line connecting the center of the start circle and the target and the line connecting the start of the movement and the hand position at peak tangential velocity.

### Model Simulations

We used the explicit cues and perturbation sequence presented across trials to simulate predictions of the COIN model according to our experimental design in all of our experiments. We ran COIN model simulations for our experiments to test whether the predictions from the COIN model matched our experimental results (See supplementary information for COIN model simulation). In our simulations, we used the original article’s default parameters (noise and model parameters)^9–11^. We also ran the Contextual Dual Rate State Space model to test whether adding contexts to the individual processes will lead to tagging and recall of distinct memories (See supplementary information for dual-rate model simulation).

### Transitional Probability

We calculated the transition probabilities between all contextual states based on the COIN simulation. In Experiments 1a and 2a, which had identical learning schedules, context A was substantially stabilized while context B was learned only briefly. This schedule resulted in minimal transition probabilities between contexts, indicating that the model predominantly maintained its current contextual state without significant shifting (Fig. 1i). Similarly, in Experiments 1b and 2b, where both contexts A and B underwent stabilized learning, the model exhibited low transition probabilities between contexts throughout the learning period (Fig. 1j). In contrast, Experiments 3a and 3b utilized an interleaved training schedule. This random perturbation during learning led to significantly higher transition probabilities between contexts, reflecting the model’s increased tendency to switch states, resulting in a higher global transition probability (Fig. 1k).

### Bias quantification

We computed a context-specific bias for every participant to assess whether the stability and recency generated any biases during expression. For each cue (Cue A, Cue B, and the No cue), we first obtained a pre-clamp “peak” (𝑃𝑒𝑎𝑘_*c*_) by averaging the motor output of the last five learning trials separately from Task A and Task B. We then averaged the motor output during the clamp phase to obtain the expression. Two time windows were used: for “early bias” - the mean of the first five clamp trials, capturing the initial, most rapid expression for the corresponding cue, was used, while the mean across the entire clamp block was used to calculate the total expression for each cue.

For each cue *c* (A, B, No cue), the fractional retention (𝑑_c_) was

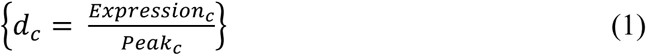

where *d_c_* = 1 indicates perfect complete contextual expression and dc = 0, no expression. We then collapsed these cue-wise measures into two orthogonal components that summarize the contextual bias (𝑑_*AB*_, only concerning cue A and cue B) and Net bias (𝐵) when relative expression (fractional retention 𝑑_*N*_) over no-cue trials was included:

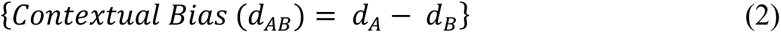

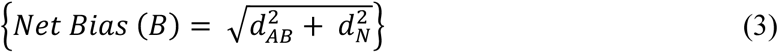

Both early and total expressions were computed per participant, and then the bias factor was calculated for each experiment separately for early and total trials. This dual-window approach allowed us to distinguish rapid context-specific biases that emerge immediately on clamp entry from the cumulative bias that accrues over the whole expression period.

## Results

To test the hypothesis that internal models of context transitions guide memory retrieval, we conducted a series of visuomotor adaptation experiments. We independently manipulated the presence of explicit contextual cues and the statistical structure of transitions between two opposing perturbations. Our results demonstrate that the expression of learned motor memories is not determined by cues alone but is powerfully shaped by learned transition statistics-specifically, the training environment’s stability and transitional entropy.

### Expression is contextual but biased by memory stability and recency

The first set of experiments examined whether contextual cues allow selective expression of motor memories and whether the stability or recency of prior experience modulates this expression. In Experiment 1a, participants first adapted to Task A (160 trials), followed by brief exposure to Task B (20 trials), each tagged with distinct contextual cues (Cue A and Cue B). This training schedule has a low transitional probability of changing context and more stability in learning A than B. Learning was similar for task A (F_[2,33]_ = 2.68, p = 0.083) and task B (F_[2,33]_ = 0.62, p = 0.541) across the three groups. In the subsequent error-clamp phase, participants were presented with one of the three contextual conditions: Cue A, Cue B, or No cue (Figure 2a). Despite Task B being learned for a short duration, the expression was still context-dependent; participants generated distinct motor outputs based on the cue presented, indicating successful separation of memories. A two-way ANOVA confirmed a main effect of the cue on expression (F_[2,33]_ = 5.89, p = 0.008, w^2^ = 0.20). Post-hoc tests revealed significant differences between Cue A and both Cue B (p=0.017) and No cue (p=0.017), although Cue B and No cue were not significantly different (p=0.928). However, analysis of the early phase of expression (first five trials) showed that even Cue B trials elicited stronger output than No cue (t_(22)_ = -2.37, p=0.027, Cohen’s d = -0.97), revealing a short-lived contextual retrieval of Task B memory despite its instability. Since later clamp trials may become contaminated by the decay of memory traces and by collapsing beliefs about context identity, particularly in the absence of ongoing error signals, we focused our analyses on the early phase of expression (first five trials), where contextual inference is most likely to reflect prior learning and not subsequent re-averaging.

**Figure 2:**
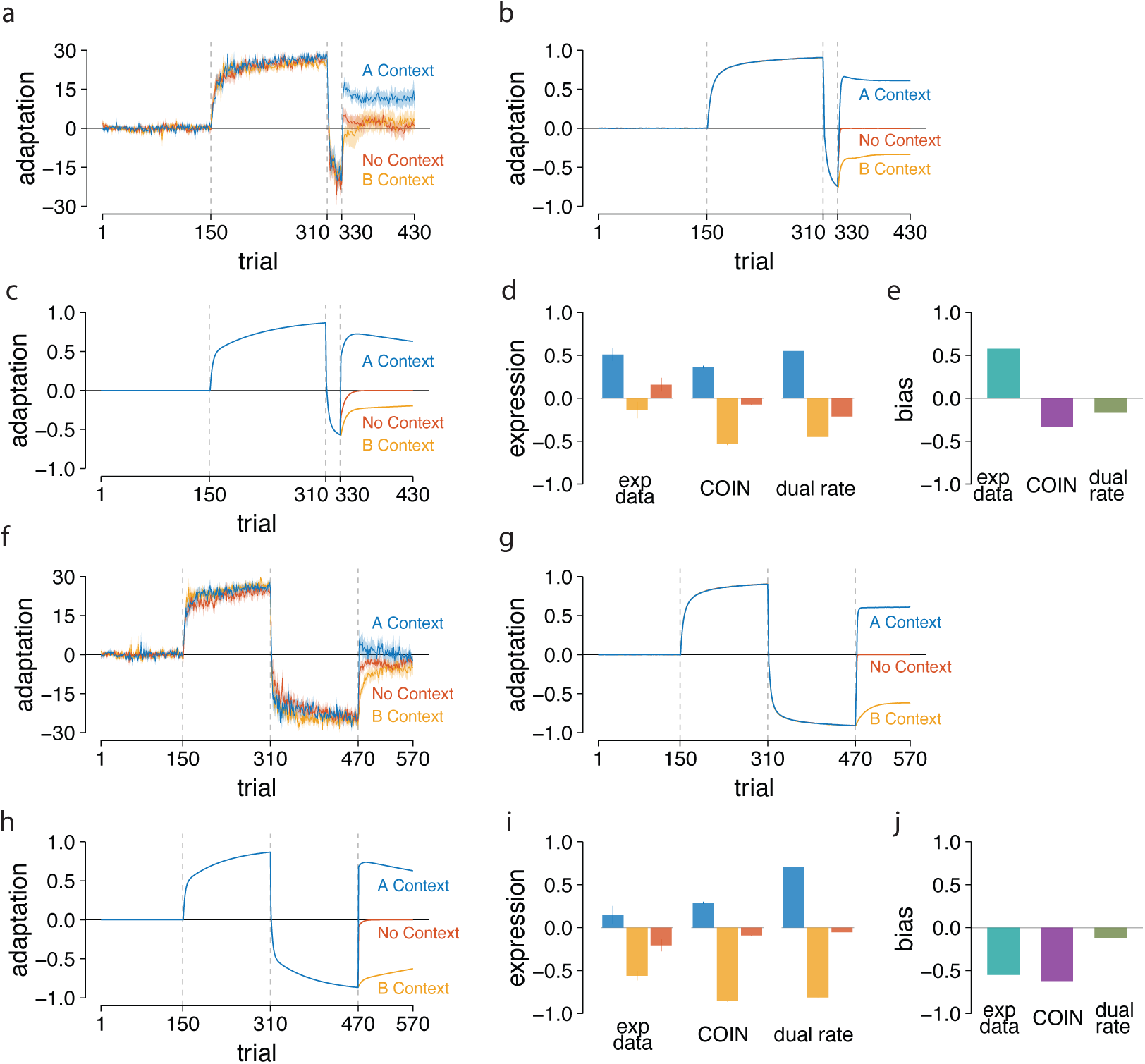
Memory expression is guided by contextual cues but biased by the stability and recency of competing memories. (a) Adaptation to the visuomotor perturbation in Experiment 1a under three contextual cues (cue A is blue, cue B is yellow, and the no cue condition is red). (b) Adaptation output from the COIN model and the dual-rate model (c). (d) Expression in the early phase (5 trials) of the Task N session of Experiment 1a. Both experimental data and model simulations show distinct memories guided by the cues. (e) Expression bias. Experimental data show recency bias; however, the COIN (purple bar) and dual-rate (green bar) models failed to predict it. (f) Adaptation output when both Task A and Task B were learned for an equal number of trials. Simulation output from the COIN model (g) and the dual-rate model (h). (i) Motor output in the early phase of the expression session of Experiment 1b. Similar to Experiment 1a, participants showed distinct memory for each contextual cue; however, the expression was biased toward the most recent memory of Task B (j). The COIN model predicted the recency bias; however, the dual-rate model failed.

Importantly, expression was not symmetric across contexts (Figure 2d). Cue A’s motor output was stronger than Cue B’s, suggesting a bias toward the more stable Task A memory. This stability-driven bias was consistent across all cue conditions, even when no cue was presented, where expression drifted toward Task A. Our bias analysis also captured this asymmetry (Figure 2e, Early bias = 0.57). The COIN and contextual dual-rate models reproduced this expression pattern (Figure 2b, 2c), with the former attributing it to the probabilistic weighting of context stability during inference. The dual-rate contextual model also produced a pattern resembling behavior (Figure 2c) because each cue gates its slow process; cue A expressed the well-trained Task A memory, while cue B retrieved the briefly trained Task B slow state. The no-cue condition expressed the baseline slow state, yielding minimal output. Bias based on the overall expression block of the COIN model simulation data indicated a positive bias towards A (bias = 0.17). Similarly, the dual-rate model predicted an overall positive bias towards A (bias = 0.36). However, both the dual-rate model (bias = -0.17) and the COIN model (bias = -0.33) predicted a negative bias during the early part of the expression (Figure 2e).

In Experiment 1b, the design was identical to 1a, except Task B was now trained for 160 trials, making both Task A and B equally stable. This schedule was designed to test the role of recency when both memories are stable, and context has a low transition probability. The learning was similar in all three groups for both task A (F_[2,33]_ = 1.05, p = 0.363) and task B (F_[2,33]_ = 0.11, p = 0.897). Participants again expressed context-specific behavior during the clamp phase (Figure 2f), with a significant main effect of cue (F_[2,33]_ = 20.72, p < 0.001, w2 = 0.52). Post-hoc tests revealed significant differences between Cue A and both Cue B (p_corrected_ <0.001) and No cue (p_corrected_ =0.005). Expression in cue B was also significantly different from cue N (p_corrected_ = 0.005). However, unlike 1a, the expression bias was reversed; motor output was stronger for Cue B than for Cue A or No cue, demonstrating a clear recency bias favoring the more recently learned Task B memory (Figure 2j). We also observed a bias towards the more recent memory B across the three groups (bias = -0.56). While now, expression of memory A with cue A was lesser than expression in experiment 1a (t_(22)_ = 2.87, p=0.008, Cohen’s d = 1.17), expression of memory B with cue B was way higher as compared to the same expression condition in experiment 1a (t_(22)_ = 3.84, p<0.001, Cohen’s d = 1.57) (Figure 2i). This pattern supports the COIN model’s prediction that recency becomes the dominant factor in determining which memory is inferred to be active when contextual stability is matched. Bias analysis confirmed this recency-driven shift (bias = - 0.62) (Figure 2j), and the COIN model captured this inversion by assigning higher posterior probability to the more recently experienced context, even under identical cue presentation (Figure 2g). Overall, COIN simulations mirrored our experimental findings, qualitatively confirming our hypothesis. On the other hand, the dual-rate model, which updates each cue’s slow process independently (Figure 2h), produced symmetry in the cue-specific expressions, and there was minimal recency bias in expression (bias = -0.12) (Figure 2j).

### Distinct contextual memories fail to drive expression in high-entropy environments

While Experiments 1a and 1b showed that contextual cues could successfully gate memory expression in stable environments, we next asked whether those memories would be expressed distinctly if the cueing environment during the test became highly volatile and dynamic. In Experiment 2a, participants followed the same acquisition schedule as in Experiment 1a (extended training on Task A: 160 trials, followed by shorter training on Task B: 20 trials). This created a stable, low-volatility environment where the transitional probability of a context change was low. However, during the testing phase, the context was switched unpredictably on a trial-by-trial basis (Figure 3a), creating a volatile, high-entropy environment. This mismatch between the stable statistics of training and the volatile statistics of testing was the key manipulation. Participants showed similar magnitude of learning as in 1a (A learning: F_[3,44]_ = 1.11, p = 0.355, B learning: F_[3,44]_ = 0.43, p = 0.729). Although overall expression showed significant cue differences (F_[2,22]_ = 12.89, p <0.001, w^2^ = 0.18), all three cue types evoked behavior in the direction of Task A, including Cue B and No Cue trials. Expression in cue A trials was significantly higher than in cue B (p_corrected_ = 0.009) or no cue trials (p_corrected_ = 0.009). There was no significant difference between expression under cue B and no cue (p_corrected_ = 0.075). Notably, the sign of early expression under Cue B flipped relative to 1a (Figure 3d), indicating that the Task B memory was not reliably retrieved despite being learned under a distinct cue. Instead, expression under all cues converged toward the more stable Task A, suggesting that participants relied on memory strength rather than cue identity to drive inference in the dynamic context. This reflects a failure of contextual gating: while the learned memories were distinct, the testing environment did not support their selective expression based on the cue.

**Figure 3:**
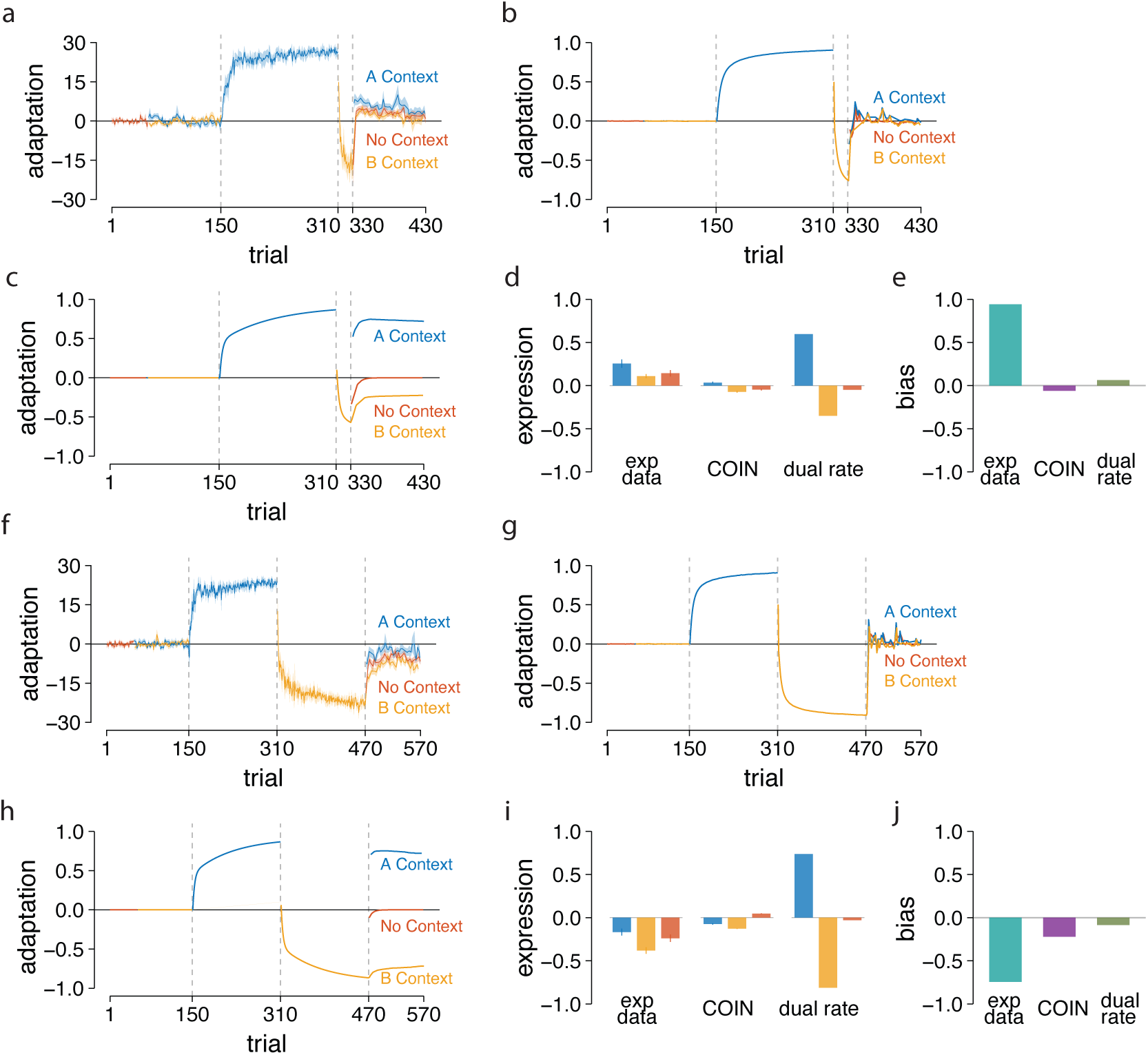
Distinct contextual cues fail to drive expression in dynamic environments, which are biased by the stability and recency of competing memories. (a) Adaptation to the visuomotor perturbation in Experiment 2a under three contextual cues (cue A is blue, cue B is yellow, and the no cue condition is red). (b) Adaptation output from the COIN model predicted the experimental data (c) and the dual-rate model failed to predict the data. (d) Expression in the early phase (5 trials) of the Task N session of Experiment 2a. Experimental data shows bias to the stable memories, and model simulations show distinct memories. (e) Expression bias. Experimental data show less recency bias than Experiment 1a; however, the COIN (purple bar) and dual-rate (green bar) models failed to predict it. (f) Adaptation output when both Task A and Task B were learned for an equal number of trials. Simulation output from the COIN model (g) and the dual-rate model (h). (i) Motor output in the early phase of the expression session of Experiment 1b. Similar to Experiment 1a, participants showed distinct memory for each contextual cue; however, the expression was biased toward the most recent memory of Task B (j). The COIN model predicted the recency bias; however, the dual-rate model failed.

To assess the interference, we compared the expression with the performance of experiment 1a, where only one cue was provided. We have found that expression in both cues A (t_(22)_ = 2.91, p = 0.008, Cohen’s d = 1.19) and B (t_(22)_ = -2.48, p = 0.021, Cohen’s d = -1.01) is reduced when cues were presented randomly. There was no significant difference between no cue expression across both experiments (t_(22)_ = 0.18, p = 0.859, Cohen’s d = 0.07). Notably, there was a greater bias towards more stable memory (early bias = 0.94) than in experiment 1a (Figure 3e).

In Experiment 2b, we asked whether this failure was due to the weak encoding of Task B. Here, both Task A and B were trained for 160 trials (as in 1b), and test cues were again interleaved. Learning of task A and task B was similar to that in experiment 1b (A learning: F_[3,44]_ = 2.10, p = 0.113, B learning: F_[3,44]_ = 0.89, p = 0.454) (Figure 3f). Similar to experiment 2a, expression differed across cues (F_[2,22]_ = 12.37, p <0.001, w^2^ = 0.27). Still, in contrast to 1b, all cue types now evoked behavior in the direction of Task B. Expression in cue B trials was significantly higher than that in cue A (p_corrected_ = 0.002) or no cue trials (p_corrected_ = 0.005). There was no significant difference between expression under cue A and no cue (p_corrected_ = 0.153).

The sign of early expression under Cue A flipped relative to 1b (Figure 3i), and even No Cue trials aligned with Task B, indicating that recency, and not cue identity, governed retrieval. We have found that expression in cue A (t_(22)_ = 2.89, p=0.008, Cohen’s d = 1.18) was flipped and got reduced in cue B (t_(22)_ = -2.70, p = 0.012, Cohen’s d = -1.10) when cues were presented randomly in comparison to Experiment 1b where single cue was presented. There was no significant difference between no cue expression across both experiments (t_(22)_ = 0.40, p = 0.69, Cohen’s d = 0.16). Notably, there was a greater bias towards more stable memory (Early bias = -0.74) than observed in experiment 1b. Thus, while memories were expressed in a consistent direction, this direction did not depend on the cue but on which memory was more recently experienced (Figure 3j). Notably, while expression in both experiments was biased toward a single memory, Task A in 2a and Task B in 2b, the magnitude of that expression still differed across cues. In Experiment 2a, although all three cues evoked Task A-like outputs, Cue A trials produced stronger expression than Cue B or No cue trials. A similar pattern was held in Experiment 2b, where Cue B trials evoked more pronounced Task B behavior than the other conditions. This suggests that while cue-memory associations were not strong enough to fully gate expression direction, they still modulated the inferred probability of each memory, consistent with partial cue-based inference. These results reinforce that context recall was degraded but not completely absent in dynamic environments.

Experiments 2a and 2b reveal a critical dissociation: although contextual memories can be acquired under stable cue-perturbation mappings, their selective expression fails when the retrieval environment is dynamic and there is a mismatch between the stable statistics of training and the volatile statistics of testing. In such cases, behavior is biased by stability or recency but not contextual, a subtle but important distinction. Participants always expressed one of the two learned memories, but this choice was dominated by stability (in 2a, Bias = 0.94 towards A, Figure 3e) or recency (in 2b, Bias = -0.74 towards B, Figure 3j), regardless of the cue presented.

Interestingly, COIN simulations did not reproduce this partial retrieval. The model predicted complete interference in both 2a (Figure 3b) and 2b (Figure 3g), with all cue conditions producing baseline-level output (Figure 3e, 3j). It also failed to capture the stability bias in 2a, which was minimal in the simulations (bias = -0.06). However, the recency bias in 2b was present (-0.22) but very short-lived. The bias based on the total expression block was minimal (-0.02) towards B. This discrepancy likely reflects a limitation in the COIN model’s current implementation, where inference at the test is tightly governed by context-transition statistics experienced during learning. Because transitions were rare in the acquisition blocks, the model expected stable contexts during expression and could not flexibly shift beliefs in a dynamic test phase.

By contrast, the contextual dual-rate model, which deterministically maps each cue to a distinct memory state, predicted strong cue-specific expression in both experiments (Figure 3c, 3h) and minimal bias (experiment 2a, bias = 0.06; experiment 2b, bias = -0.09) (Figure 3e, 3j), again failing to capture the observed interference. Neither model could explain the pattern of non-contextual but biased expression seen in human participants, suggesting the need for future model refinements that allow for more flexible inference under changing contextual priors.

### Learning in a high transitional probability environment leads to more flexible expression

The interference observed in Experiments 2a and 2b, but not in Experiments 1a and 1b, prompted us to ask whether flexibility in expression depends on the nature of the environment in which memories were acquired. Specifically, we hypothesized that learning in an environment with high context-transition probabilities might promote more adaptable inference during tests, enabling selective expression even under dynamically changing cues.

In Experiment 3a, participants learned Task A and Task B under a dynamic training environment in which cues (Cue A, Cue B, No Cue) were randomly interleaved. Crucially, they experienced an equal number of trials for both tasks (Figure 4a). Participants were able to adapt to both A (F_[1,11]_ = 71.75, p<0.001, w^2^ = 0.74) and B (F_[1,11]_ = 81.06, p <0.001, w^2^ = 0.67) perturbations without interference. The magnitude of learning by the end of training was comparable across the two tasks (t_(11)_ = 0.93, p = 0.368). During the clamp phase, when contextual cues were again presented in a randomly interleaved manner, participants displayed clean cue-specific expression of the corresponding memories with no evidence of interference (F_[2,22]_ = 20.71, p<0.001, w^2^ = 0.56) (Figure 4d). Participants showed distinct behavior when presented with different cues (p<0.005). Importantly, there was no discernible bias in expression (-0.01), consistent with the fact that both memories were equally stable and recent (Figure 4e).

**Figure 4:**
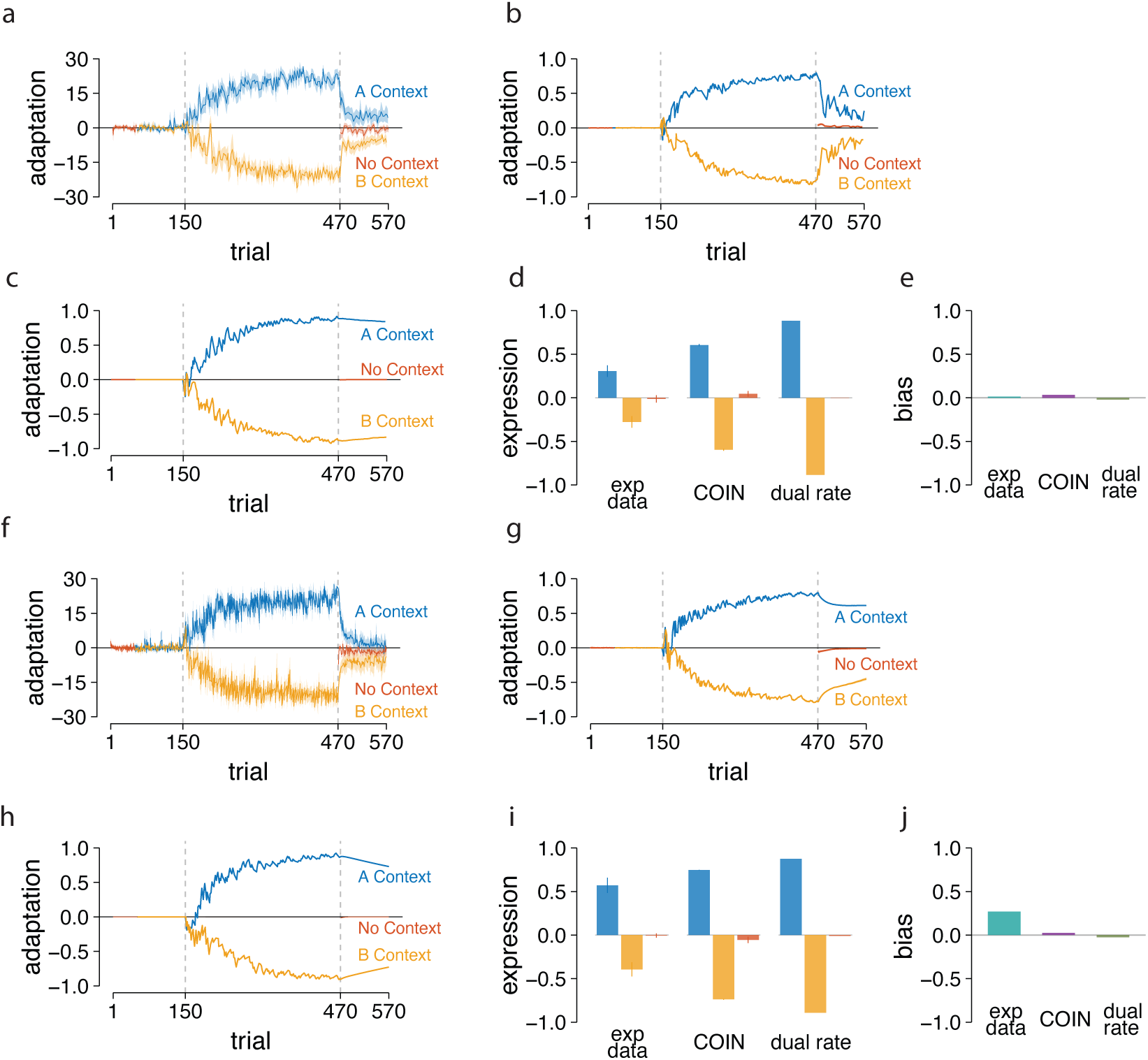
Learning in a dynamic environment leads to more flexible expression. Memory expression is more flexible when learnt in a dynamic environment and minimally biased by competeing memories. (a) Adaptation to the visuomotor perturbation in Experiment 3a under three contextual cues learned and expressed randomly (cue A is blue, cue B is yellow, and the no cue condition is red). (b) Adaptation output from the COIN model and (c) the dual-rate model. (d) Expression in the early phase (5 trials) of the Task N session of Experiment 3a. Both experimental data and model simulations show distinct memories guided by the cues. (e) Expression bias. Experimental data show minimal bias and were cue dependent; the COIN (purple bar) and dual-rate (green bar) models also predict it. (f) Adaptation output in Experiment 3b under three contextual cues learned randomly, and expressed only one cue per participant. Simulation output from the COIN model (g) and the dual-rate model (h). (i) Motor output in the early phase of the expression session of Experiment 3b. Similar to Experiment 3a, participants showed distinct memory for each contextual cue; Expression was cue dependent (j). The COIN model and the dual-rate model predicted the same expression.

These results suggest that learning under high transition probabilities facilitates the creation of cue-tagged memories that can be selectively recalled even when test cues are interleaved. Both the COIN model and the contextual dual-rate model correctly predicted this outcome. In the COIN model, the matched structure between learning and test (i.e., dynamic cue switching in both phases) sustained the reliability of the cues, enabling accurate context inference (Figure 4b). Likewise, the dual-rate model produced the expected result because each cue consistently activated its corresponding slow process (Figure 4c). Both models also showed no bias towards any specific memory (COIN: -0.03; dual-rate: -0.02, Figure 4e).

We next asked whether memories acquired in a dynamic environment could also support selective expression when tested in a stable environment. In Experiment 3b, three separate groups learned Tasks A and B under interleaved conditions, but the test phase presented only one contextual cue per group (Cue A, Cue B, or No Cue) (Figure 4f). For cue A, participants showed improvement over the course of training (F_[1,33]_ = 237.14, p < 0.001, w^2^ = 0.72), and there was no significant difference across groups (F_[2,33]_ = 0.85, p =0.438). Similarly, for cue B, participants reduced the error over the training period (F_[1,33]_ = 149.68, p < 0.001, w^2^ = 0.69), and all the groups were similar in the magnitude of learning (F_[2,33]_ = 0.85, p =0.435). Participants showed a similar learning magnitude for tasks A and B (F_[2,33]_ = 0.72, p = 0.494). In the clamp phase, each group expressed the memory corresponding to the cue they received (F_[2,33]_ = 47.75, p<0.001, w^2^ = 0.72), indicating no interference (Figure 4i). Participants showed distinct behavior when presented with different cues (p<0.001). Similar to Experiment 3a, there was minimal bias in expression (-0.27), suggesting that both memories were equally stable and recent (Figure 4j).

These findings provide further support for the idea that learning in a high transitional probability environment builds more flexible and robust context-memory associations. Both COIN and dual-rate models replicated this result: COIN because prior exposure to cue switching enabled the inference of dynamic contextual states (Figure 4g), and the dual-rate model due to its structural mapping of cue to slow state (Figure 4h). Similar to the actual behavior, both models also showed minimal bias (COIN: 0.02; dual-rate: -0.02)

Together, Experiments 3a and 3b demonstrate that training in dynamic environments enables more flexible and context-appropriate memory expression, regardless of whether the testing environment is stable or dynamic. This supports a central prediction of the COIN model: the motor system incorporates the history of contextual cues and their transition statistics into its inference about which memory to express.

## Discussion

Our study provides direct behavioral evidence that transition statistics - the learned temporal structure of context switches - shape memory retrieval. Across six visuomotor adaptation experiments, we manipulated training order, stability, and entropy to test how memories of opposing perturbations are recalled under uncertainty. The results reveal that retrieval is not governed by cues alone, but by an arbitration process in which cues are weighted against priors derived from transition statistics. This principle, while demonstrated here in motor learning, extends broadly to memory systems, suggesting a domain-general mechanism of context-sensitive retrieval. These findings align with classic ideas of hippocampal indexing^27^, predictive coding models of memory^28,29^, and information-theoretic principles of surprise and uncertainty, situating our results within a broader theoretical landscape^30–32^.

### Stability, Recency, and Volatility

Blocked training established strong priors for continuity. In Experiment 1a, where one memory (Task A) was practiced extensively and the other (Task B) only briefly (∼20 trials), retrieval largely defaulted to the stable memory - the one with greater cumulative practice - even when cues signaled the alternative mapping. This indicates that even minimal exposure to Task B was sufficient to form a distinct, context-linked memory, but its expression was overshadowed by the stronger stability prior. In Experiment 1b, where stabilities were matched, retrieval was biased toward the more recent memory, underscoring that in the absence of a stability advantage, recency dominates. These patterns show that retrieval biases are lawful consequences of learned transition statistics: stability priors weight older, more experienced memories, while volatility priors bias toward more recent ones. Such biases mirror those in the broader memory literature. For instance, the recency effect - recent items being readily recalled due to lingering availability in short-term memory^33,34^ - parallels the dominance of recent memories in volatile or equal-stability contexts. Conversely, the primacy effect (favoring early, well-practiced items) is analogous to stability priors weighting older memories^34^. Additionally, stability biases in metamemory lead people to overestimate the durability of memories (underestimating forgetting), aligning with the persistent influence of stable memories in our experiments^35^.

By contrast, interleaved training (Experiments 3a and 3b) cultivated high-entropy priors that preserved cue-specific memory separation even in volatile test conditions. Retrieval remained highly flexible, with minimal stability or recency bias, because participants had learned to expect frequent context switches. This suggests that interleaved exposure induces a meta-learning of volatility - effectively training the learner’s inference policies to maintain multiple competing memories in readiness and to arbitrate retrieval flexibly based on context cues.

### Arbitration Between Cues and Transition Priors

Together, our findings converge on a single principle: retrieval is an arbitration between cue information and transition priors. In stable environments, priors favor continuity, yielding persistence or stability biases. When stabilities are equal, priors favor recency, yielding recency biases. When training and testing structures mismatch, priors dominate and cue-based gating collapses, producing biased but non-contextual behavior. Conversely, when volatility is high, priors flatten, allowing cues to regain control of behavior.

Empirically, when training and test structures mismatched (Experiments 2a and 2b), cue-based retrieval collapsed. Despite valid cues at test, participants defaulted almost entirely to the stable memory (Exp 2a) or the recent memory (Exp 2b), with contextual cues modulating only the *magnitude* of responses, not the direction. This demonstrates that retrieval failure arises not from an inability to encode or recall cue associations, but from a mismatch between the learned transition structure and the test environment. It reframes anterograde interference not as an encoding deficit, but as a problem of inference: strong stability or recency priors from earlier training overwhelm contradictory cue evidence when the context suddenly becomes unpredictable. In other words, what traditionally has been described as failure to learn or express a new memory due to prior learning can be recast as an arbitration issue - where robust transition priors from a stable or recent history suppress the expression of newly acquired mappings (even when cues indicate a switch), resulting in inference-based “blocking” rather than true learning failure.

This arbitration framework also helps reinterpret spontaneous recovery - where a dormant memory re-emerges after a delay or context change - as a shift in inferred transition statistics. When the environment appears stable, the system reweights older, more practiced memories with high stability priors, allowing a previously suppressed memory to resurface. Similarly, anterograde interference reflects the converse: when the context is perceived as continuous or when recent practice dominates, strong continuity or recency priors hinder the retrieval of a newer memory. This underscores how spontaneous recovery and anterograde interference phenomena can be explained by models that incorporate transition statistics to adjust memory expression under changing environmental structures. Likewise, partial retrieval effects - where cues influence the *extent* of expression, but not which memory is expressed (as seen in our volatile test conditions) - reflect a probabilistic weighting of multiple memories rather than an all-or-none gating. These ideas mirror latent-state inference mechanisms in memory and perception, and align with recent Bayesian models like the Contextual Inference (COIN) model^9–11^ that jointly infer context identity and its dynamics.

The COIN model’s Bayesian framework provides a natural account of such arbitration through posterior inference: P(context|cue, prior) ∝ P(cue|context) × P(context|prior). Although the COIN model framework explains that the stability effects emerge through Bayesian weighting of transition probabilities, where high self-transition probabilities (κ ≈ 0.7) favor persistent memories. However, our simulations revealed that COIN’s default implementation fails to capture the partial retrieval patterns in Exp 2. The model’s fixed transition assumptions predict either full interference or complete separation, missing the graded bias effects we observed. This failure points to a crucial limitation: COIN assumes that training-derived priors remain static during testing. Recent extensions exploring probabilistic context inference suggest that dynamic updating of transition beliefs may be necessary to account for the full range of adaptation phenomena. Specifically, parameter adjustments (e.g., increasing volatility sensitivity α from ∼0.1 to ∼0.3) or incorporating meta-learning of transition statistics could enable the model to capture our data. Such extensions would allow the system to detect mismatches between expected and observed volatility, modulating the strength of prior-driven biases accordingly.

In contrast, simple dual-rate state-space models^36^ can approximate the empirical stability and recency biases via fast and slow learning processes, but they lack explicit representation of context and transition statistics - limiting their explanatory power. Our data suggest that retrieval phenomena such as contextual interference, cue-induced collapse, and spontaneous recovery require models that explicitly represent transition priors.

### Cross-Domain Parallels and Cognitive Theory

The logic of transition-guided retrieval appears to generalize beyond motor learning. In episodic memory, for example, event boundaries (i.e. abrupt increases in volatility) segment experiences into distinct contexts and reduce interference between events^14,21^. In working memory, recall sequences naturally cluster according to learned transition probabilities^23^, paralleling the recency-weighted arbitration we observed in motor adaptation. In language acquisition, infants extract word boundaries from statistical transitions in syllables^18^, and bilinguals with extensive switching experience show reduced switch costs^19^ - akin to our interleaved-training participants who learned to switch seamlessly. Even in perception and decision-making, adaptation slows in volatile environments^8,22^, consistent with a dampening of cue-driven updating when transitions are unpredictable.

Our results also reinforce principles like event segmentation theory^37^ and the classic encoding-specificity principle^38,39^. When transitions are predictable and stable, cue–context associations are strengthened and memory retrieval is more straightforward. When environmental entropy rises, cue efficacy declines as the system places more weight on recent experience. In essence, the brain encodes not only *what* cues occur, but *how reliably* they occur in sequence. This enables context-sensitive retrieval guided by latent statistical structure: memory recall is tuned not just to the presence of a cue, but to the learned reliability of that cue given the history of context switches.

### Neural Implementation: Precision Gating and Memory Arbitration

The interaction between contextual cues and transition statistics is likely supported by a network involving the prefrontal cortex (PFC), basal ganglia, and cerebellum, which collectively learn transitional probabilities to gate motor memory expression in the primary motor cortex (M1). The PFC is hypothesized to maintain and flexibly apply contextual priors^28^, the basal ganglia learns state transition statistics through reinforcement mechanisms^40^, and the cerebellum fine-tunes the predictive timing of these transitions^41^. These brain areas then bias competition between memory representations in M1, which are thought to be stored in distinct neural subspaces^25^. Our findings - that expression is governed by learned stability and recency and collapses under statistical mismatch-support this framework, suggesting that these fronto-subcortical circuits gate memory retrieval^42^. Future studies could test these interactions using fMRI to track PFC-basal ganglia connectivity during volatility shifts, EEG to measure theta-band oscillations as a signature of context switching, or TMS to disrupt PFC or cerebellar processing and probe their causal roles in the arbitration of competing motor memories.

### Summary

By demonstrating that retrieval reflects an arbitration between cues and transition priors, our study reframes memory recall as a predictive process governed by learned environmental dynamics. This principle has direct implications across domains. For example, interleaved practice (high-context variability) in skill acquisition and coaching could promote volatility-tuned retrieval policies that generalize better across changing environments, aligning with classical findings that variable practice enhances long-term adaptability. In rehabilitation settings, manipulating contextual entropy (e.g. practicing tasks in unpredictable sequences) could help patients to regain flexible switching between motor memories, potentially reducing interference in re-learning movements.

More broadly, this work contributes to a domain-general theory of memory retrieval. It argues that memory is not a static cache indexed only by cues, but rather a dynamic inference process guided by the brain’s internal model of temporal structure. Transition statistics - how often things change and in what patterns - become an integral part of the memory code. Whether in perception, action, language, or episodic recall, the same principle applies: the brain learns not only *what* to expect, but *when* to expect it-and uses this temporal knowledge to decide which memory to recall in a given moment.

## ACKNOWLEDGEMENT

We thank IIT Hyderabad for all the institutional-level support.

## References

1. Caithness, G. et al. Failure to Consolidate the Consolidation Theory of Learning for Sensorimotor Adaptation Tasks. J. Neurosci. 24, 8662–8671 (2004).

2. Krakauer, J. W., Ghilardi, M. F. & Ghez, C. Independent learning of internal models for kinematic and dynamic control of reaching. Nat. Neurosci. 2, 1026–1031 (1999).

3. Howard, I. S., Ingram, J. N., Franklin, D. W. & Wolpert, D. M. Gone in 0.6 Seconds: The Encoding of Motor Memories Depends on Recent Sensorimotor States. J. Neurosci. 32, 12756–12768 (2012).

4. Kojima, Y., Iwamoto, Y. & Yoshida, K. Memory of learning facilitates saccadic adaptation in the monkey. J. Neurosci. Off. J. Soc. Neurosci. 24, 7531–7539 (2004).

5. Sarwary, A. M. E., Stegeman, D. F., Selen, L. P. J. & Medendorp, W. P. Generalization and transfer of contextual cues in motor learning. J. Neurophysiol. 114, 1565–1576 (2015).

6. Addou, T., Krouchev, N. & Kalaska, J. F. Colored context cues can facilitate the ability to learn and to switch between multiple dynamical force fields. J. Neurophysiol. 106, 163–183 (2011).

7. Carvalho, P. F. & Goldstone, R. L. The benefits of interleaved and blocked study: Different tasks benefit from different schedules of study. Psychon. Bull. Rev. 22, 281–288 (2015).

8. Gonzalez Castro, L. N., Hadjiosif, A. M., Hemphill, M. A. & Smith, M. A. Environmental Consistency Determines the Rate of Motor Adaptation. Curr. Biol. 24, 1050–1061 (2014).

9. Heald, J. B., Lengyel, M. & Wolpert, D. M. Contextual inference underlies the learning of sensorimotor repertoires. Nature 600, 489–493 (2021).

10. Heald, J. B., Lengyel, M. & Wolpert, D. M. Contextual inference in learning and memory. Trends Cogn. Sci. 27, 43–64 (2023).

11. Heald, J. B., Wolpert, D. M. & Lengyel, M. The Computational and Neural Bases of Context-Dependent Learning. Annu. Rev. Neurosci. 46, 233–258 (2023).

12. Nissen, M. J. & Bullemer, P. Attentional requirements of learning: Evidence from performance measures. Cognit. Psychol. 19, 1–32 (1987).

13. Song, S., Howard, J. H. & Howard, D. V. Implicit probabilistic sequence learning is independent of explicit awareness. Learn. Mem. Cold Spring Harb. N 14, 167–176 (2007).

14. Clewett, D. & Davachi, L. The ebb and flow of experience determines the temporal structure of memory. Curr. Opin. Behav. Sci. 17, 186–193 (2017).

15. Radvansky, G. A. & Zacks, J. M. Event boundaries in memory and cognition. Curr. Opin. Behav. Sci. 17, 133–140 (2017).

16. Stachenfeld, K. L., Botvinick, M. M. & Gershman, S. J. The hippocampus as a predictive map. Nat. Neurosci. 20, 1643–1653 (2017).

17. Kuhl, P. K. Early language acquisition: cracking the speech code. Nat. Rev. Neurosci. 5, 831– 843 (2004).

18. Saffran, J. R., Aslin, R. N. & Newport, E. L. Statistical Learning by 8-Month-Old Infants. Science 274, 1926–1928 (1996).

19. Gullifer, J. W. & Titone, D. Engaging proactive control: Influences of diverse language experiences using insights from machine learning. J. Exp. Psychol. Gen. 150, 414–430 (2021).

20. Jiang, Y. V., Swallow, K. M., Rosenbaum, G. M. & Herzig, C. Rapid acquisition but slow extinction of an attentional bias in space. J. Exp. Psychol. Hum. Percept. Perform. 39, 87–99 (2013).

21. Nobre, A. C. & Van Ede, F. Anticipated moments: temporal structure in attention. Nat. Rev. Neurosci. 19, 34–48 (2018).

22. Behrens, T. E. J., Woolrich, M. W., Walton, M. E. & Rushworth, M. F. S. Learning the value of information in an uncertain world. Nat. Neurosci. 10, 1214–1221 (2007).

23. Polyn, S. M., Norman, K. A. & Kahana, M. J. A context maintenance and retrieval model of organizational processes in free recall. Psychol. Rev. 116, 129–156 (2009).

24. Cisek, P. & Kalaska, J. F. Neural mechanisms for interacting with a world full of action choices. Annu. Rev. Neurosci. 33, 269–298 (2010).

25. Churchland, M. M. et al. Neural population dynamics during reaching. Nature 487, 51–56 (2012).

26. Oldfield, R. C. The assessment and analysis of handedness: The Edinburgh inventory. Neuropsychologia 9, 97–113 (1971).

27. Hirsh, R. The hippocampus and contextual retrieval of information from memory: A theory. Behav. Biol. 12, 421–444 (1974).

28. Barron, H. C., Auksztulewicz, R. & Friston, K. Prediction and memory: A predictive coding account. Prog. Neurobiol. 192, 101821 (2020).

29. Friston, K. A theory of cortical responses. Philos. Trans. R. Soc. B Biol. Sci. 360, 815–836 (2005).

30. Baldi, P. & Itti, L. Of bits and wows: A Bayesian theory of surprise with applications to attention. Neural Netw. 23, 649–666 (2010).

31. Itti, L. & Baldi, P. Bayesian surprise attracts human attention. Vision Res. 49, 1295–1306 (2009).

32. Knill, D. C. & Pouget, A. The Bayesian brain: the role of uncertainty in neural coding and computation. Trends Neurosci. 27, 712–719 (2004).

33. Baddeley, A. D. & Hitch, G. Working Memory. in Psychology of Learning and Motivation vol. 8 47–89 (Elsevier, 1974).

34. Glanzer, M. & Cunitz, A. R. Two storage mechanisms in free recall. J. Verbal Learn. Verbal Behav. 5, 351–360 (1966).

35. Kornell, N. & Bjork, R. A. A stability bias in human memory: Overestimating remembering and underestimating learning. J. Exp. Psychol. Gen. 138, 449–468 (2009).

36. Smith, M. A., Ghazizadeh, A. & Shadmehr, R. Interacting adaptive processes with different timescales underlie short-term motor learning. PLoS Biol. 4, e179 (2006).

37. Zacks, J. M., Speer, N. K., Swallow, K. M., Braver, T. S. & Reynolds, J. R. Event perception: A mind-brain perspective. Psychol. Bull. 133, 273–293 (2007).

38. Tulving, E. Elements of Episodic Memory. (Clarendon Press ; Oxford University Press, Oxford [Oxfordshire]: New York, 1983).

39. Tulving, E. & Thomson, D. M. Encoding specificity and retrieval processes in episodic memory. Psychol. Rev. 80, 352–373 (1973).

40. Doya, K. What are the computations of the cerebellum, the basal ganglia and the cerebral cortex? Neural Netw. Off. J. Int. Neural Netw. Soc. 12, 961–974 (1999).

41. Ivry, R. B. & Spencer, R. M. The neural representation of time. Curr. Opin. Neurobiol. 14, 225–232 (2004).

42. Courtin, J. et al. Prefrontal parvalbumin interneurons shape neuronal activity to drive fear expression. Nature 505, 92–96 (2014).

