## Supplementary Material for "Contextual Cues and Transition Statistics Drive Expression of Competing Motor Memories"

**Supplementary Information**

*Computational method*

Conventional state space models for motor learning do not account for the representation of multiple memories and how they interact. Thus, they cannot explain how we can learn, update, and express multiple motor skills in parallel without any interference. The COIN model proposed by Heald et al. puts the repertoire of multiple motor memories at the core[1]. It solves the problem of multiple memories by proposing that each memory is tagged by a context and inference about which context is active at an instance to drive how all the memories are updated and expressed, or if a new memory would be created. We implemented this model and asked questions based on the predictions.

The COIN model posits that the environment can be represented by an infinite number of discrete contexts, each associated with a time-varying state. These contexts transition according to a Markov process. The active context at any given time determines the observed state feedback, which is influenced by the learner's motor actions. The model also incorporates sensory cues that help infer the current context. Each context is associated with a set of parameters that define the perturbation and the probability of that context being active.

The original article proposing the COIN model explains the model in detail. Here, we go through the summary of the model in the context of visuomotor adaptation:

1. Latent state dynamics

Each context j maintains a hidden state $x_t^{(j)}$, the learner's estimate of that context's rotation. Across trials, it obeys a linear–Gaussian update:

$$\left\{x^{j}_{t+1;} = a_{j,x^{j}_{t;}} + d_j + \xi_t \qquad \xi_t \sim N(0,\sigma_w^2)\right\}$$ ----- (1)

Where $a_j \in (0,1)$ is the retention factor (larger ⇒ more stable, slower decay), $d_j$ is drift, so the long-run mean $\frac{\{d_j\}}{\{1-a_j\}}$ can differ from zero, and

$\xi_t^{(j)}$ is process noise for context j, with process-noise variance $\sigma_w^2$, modeling unsignaled state fluctuations

2. Observation Model

When context $c_t$ is active, the observed cursor angle $y_t$ is the sum of its state plus a fixed bias and sensory/motor noise:

$$\left\{y_t = x_t^{\{(c_t)\}} + b_{\{c_t\}} + v_t \qquad v_t \sim N\left(0, \sigma_y^2\right)\right\} \qquad \text{----- (2)}$$

Where $b_j$ is imposed visuomotor rotation  and $v_t$ is total observation noise (motor and sensory) with variance $\sigma_y^2$.

3. Context transitions

Contexts switch according to a *sticky* HDP-HMM, with

$$\left\{P\left(c_t = k \mid c_{\{t-1\}} = j\right) = \pi_{\{jk\}}, \qquad \pi_{\{jj\}} \propto k + \alpha\beta j, ; \pi_{\{jk\neq j\}} \propto \alpha\beta_k\right\} \qquad \text{----- (3)}$$

Where α, β are global transition weights (stick-breaking), κ is self-transition bias, encouraging contextual persistence. It allows *on-line creation* of new contexts whenever none of the existing contexts explains the data well.

4. Bayesian context inference

On each trial, the model fuses the sensory cue $q_t$ and the prediction error $y_t - \hat{y}_{t|t-1}^{(j)}$ to update the posterior "responsibilities"

$$\left\{P\left(c_t = j \mid q_t, y_t, y_{\{1:t-1\}}\right) \propto P\left(y_t \mid c_t = j\right) \times P\left(c_t = j \mid q_t, y_{\{1:t-1\}}\right)\right\} \qquad \text{----- (4)}$$

This recursion blends the *likelihood* of each context generating the observed error with the *prior* from cue-based and transition-based expectations.

5. Motor command & parallel learning

Before seeing feedback on trial t, the controller issues

$$\left\{ u_t = \sum_j \rho_t^{\{(j)\}} \left[ \hat{x}^{\{(j)\}}_{\{t|t-1\}} + b_j \right], \rho_t^{\{(j)\}} = P\left( c_t = j \mid q_t, y_{\{1:t-1\}} \right) \right\} \quad \text{----- (5)}$$

so that contexts with higher responsibility $\rho_t^{(j)}$ contribute more. After feedback, every $x^{(j)}$ is updated by a scaled Kalman gain $\propto \rho_t^{(j)}$, ensuring *parallel* but *weighted* memory updating.

These equations form the core of how the COIN model works for visuomotor adaptation. The model infers the current context based on sensory cues and movement errors, updates its beliefs about the rotation associated with each context, and combines these beliefs to produce adapted movements. In context of our study, extended training would raise retention $a_j$, reducing uncertainty and boosting $\rho_t^{(j)}$ in Eq (5) for that context. When contexts have equal $a_j$, the transition prior $\pi_{jk}$ (Eq 3) favours the most recently visited context, flipping $\rho_t^{(j)}$ . Also, randomly interleaving contexts increases entropy in $\pi_{jk}$ , flattening $\rho_t^{(j)}$ and diminishing cue-based separation, producing the observed interference patterns.

*Multi-rate contextual state space model*

To determine whether contextual Bayesian inference is essential, or whether cue-specific state-space learning alone can account for our data, we simulated the contextual dual-rate state-space model initially proposed by Lee and Schweighofer (2009)[2].

This model extends the two-time-scale framework of Smith, Ghazizadeh, and Shadmehr (2006)[3] by assigning a separate slow process to every discrete cue while retaining a single, cue-independent fast process. On trial *n* the motor command is therefore the algebraic sum of the fast state $x_{f,n}$ and the slow state $x_{s,c,n}$ belonging to the currently cued context c:

$$\{y_n = x_{f,n} + x_{s,c,n}\} \quad \text{----- (6)}$$

After the movement, the learner observes a sensory-prediction error

$$\{e_n = y_n + p_n + v_n\} \quad \text{----- (7)}$$

Where $p_n$ is the imposed cursor rotation and $v_n \sim N(0,\sigma^2)$ represents motor and measurement noise. Only the slow state associated with the active cue is modified; all other slow states decay. State evolution is therefore

$$\{x_{s,c,n+1} = A_s * x_{s,c,n} - B_s * e_n\} \quad \text{----- (8)}$$

$$\{x_{s,c,n+1} = A_s * x_{s,j \neq c,n}\} \quad \text{----- (9)}$$

$$\{x_{f,n+1} = A_f * x_{f,n} - B_f * e_n\} \quad \text{----- (10)}$$

A is retention factor, B is learning rate, s is slow, f is fast.
with parameters taken directly from Lee and Schweighofer's fits: $A_s = 0.998$, $A_f = 0.85$, $B_s = 0.021$, $B_f = 0.11$, and $\sigma = 0.001$.
Because no probability weighting is applied, the cue presented on a given trial fully determines which slow state contributes to behavior. In contrast with the COIN model, there is no "mixing" of memories; interference is prevented simply because the relevant slow state is switched off whenever its cue is absent.

**References**

1. Heald, J. B., Lengyel, M. & Wolpert, D. M. Contextual inference underlies the learning of sensorimotor repertoires. *Nature* **600**, 489–493 (2021).

2. Lee, J.-Y. & Schweighofer, N. Dual adaptation supports a parallel architecture of motor memory. *J. Neurosci. Off. J. Soc. Neurosci.* **29**, 10396–10404 (2009).

3. Smith, M. A., Ghazizadeh, A. & Shadmehr, R. Interacting adaptive processes with different timescales underlie short-term motor learning. *PLoS Biol.* **4**, e179 (2006).
